# Herpes Simplex Virus 1 (HSV-1) Uses the Rab6 Post-Golgi Secretory Pathway For Viral Egress

**DOI:** 10.1101/2023.12.13.571414

**Authors:** Melissa H. Bergeman, Kimberly Velarde, Honor L. Glenn, Ian B. Hogue

## Abstract

Herpes Simplex Virus 1 (HSV-1) is an alpha herpesvirus that infects a majority of the world population. The mechanisms and cellular host factors involved in the intracellular transport and exocytosis of HSV-1 particles are not fully understood. To elucidate these late steps in the replication cycle, we developed a live-cell fluorescence microscopy assay of HSV-1 virion intracellular trafficking and exocytosis. This method allows us to track individual virus particles, and identify the precise moment and location of particle exocytosis using a pH-sensitive reporter. We show that HSV-1 uses the host Rab6 post-Golgi secretory pathway during egress. The small GTPase, Rab6, binds to nascent secretory vesicles at the *trans*-Golgi network and regulates vesicle trafficking and exocytosis at the plasma membrane. HSV-1 particles colocalize with Rab6a in the region of the Golgi, cotraffic with Rab6a to the cell periphery, and undergo exocytosis from Rab6a vesicles. Consistent with previous reports, we find that HSV-1 particles accumulate at preferential egress sites in infected cells. The Rab6a secretory pathway mediates this preferential/polarized egress, since Rab6a vesicles accumulate near the plasma membrane similarly in uninfected cells. These data suggest that, following particle envelopment, HSV-1 egress follows a pre-existing cellular secretory pathway to exit infected cells rather than novel, virus-induced mechanisms.

**IMPORTANCE:** HSV-1 infects a majority of people. It establishes a life-long latent infection, and occasionally reactivates, typically causing characteristic oral or genital lesions. Rarely in healthy natural hosts, but more commonly in zoonotic infections and in elderly, newborn, or immunocompromised patients, HSV-1 can cause severe herpes encephalitis. The precise cellular mechanisms used by HSV-1 remain an important area of research. In particular, the egress pathways that newly-assembled virus particles use to exit from infected cells are unclear. In this study, we used fluorescence microscopy to visualize individual virus particles exiting from cells, and found that HSV-1 particles use the pre-existing cellular secretory pathway, regulated by the cellular protein, Rab6.

## INTRODUCTION

During the late stages of viral replication, HSV-1 particles traffic to the plasma membrane to undergo exocytosis and complete the infectious cycle. The details of this process, including which host factors are associated, and whether HSV-1 uses pre-existing cellular pathways or induces novel virus-induced pathways, are little understood. Here we describe how HSV-1 particles use the Rab6a secretory pathway to traffic from the region of the Golgi to the plasma membrane for viral egress by exocytosis.

A member of the Ras GTPase superfamily, Rab6a is a highly conserved protein across eukaryotic organisms [1]. As molecular switches, Rab proteins alternate between an active form that binds GTP and an inactive form that binds GDP. In their GTP-bound active form, Rab proteins localize to particular intracellular membranes, and function to recruit a wide variety of effector proteins that determine organelle identity and direct vesicular traffic between organelles [1]. Rab6a localizes strongly to the Golgi and is involved in ER-Golgi and intra-Golgi vesicular traffic. Importantly, Rab6a is present on nascent secretory vesicles that bud from the Trans-Golgi Network (TGN), and directs their intracellular transport and exocytosis at the plasma membrane [2, 3, 4, 5].

Rab6a has been identified as an important host factor for replication and intracellular transport/egress of several types of viruses. Parvovirus capsids have been shown to associate with Rab6a secretory vesicles [6], and genetic screens have shown that HIV requires Rab6a for infection [7]. The gamma herpesvirus human cytomegalovirus (HCMV) requires Rab6a for viral protein trafficking to the viral assembly compartment [8, 9]. Alpha herpesviruses Varicella Zoster Virus (VZV) and Pseudorabies Virus (PRV) are associated with Rab6a, and PRV undergoes exocytosis from Rab6a secretory vesicles [10, 11, 12]. In HSV-1, Rab6 contributes to proper intracellular trafficking of viral membrane proteins prior to secondary envelopment [13, 14], but the role of Rab6 in post-assembly trafficking and egress is not known.

Using a range of imaging modalities for both fixed and live cells, we show that HSV-1 progeny virus particles colocalize with Rab6a, cotransport with Rab6a from the Golgi region to the cell periphery, and complete exocytosis from Rab6a secretory vesicles. In addition, we show that Rab6a preferentially accumulates at particular locations at the adherent corners, cell extensions, and cell-cell junctions, in both infected and uninfected cells. Our data suggest that HSV-1 egress uses pre-existing cellular pathways, rather than inducing novel viral-specific trafficking routes.

## RESULTS

### Rab6a and HSV-1 Capsid Colocalize to the *trans*-Golgi Region

Rab6 has a well-established role as a marker of the Golgi apparatus [2, 3, 5]. Rab6 is considered one of the few *bona fide* Golgi Rab proteins, whereas others, like Rab8 and Rab11 participate in both Golgi and endocytic trafficking[15, 16]. Thus, we performed fluorescent superresolution microscopy on cells that had been transduced to express a fluorescent Rab6a protein and were also coinfected with a fluorescent reporter strain of HSV-1. We transduced Vero (African green monkey kidney) cells with an HSV-1-based amplicon vector that expresses an EmGFP-Rab6a transgene. The amplicon vector consists of a bacterial plasmid, transgene expression cassette, and origin of replication and packaging signal sequences from the HSV-1 genome. In the presence of a helper virus, the amplicon vector is replicated and packaged into HSV-1 particles, producing a mixed virus stock containing both amplicon and helper virus genomes. We used HSV-1 OK14 [17], based on the 17syn^+^ laboratory strain, as the helper virus because it expresses an mRFP-VP26 capsid tag to image virus particles in infected cells. Vero cells were fixed and probed for the *trans*-Golgi marker Golgin97 at 6 hours post infection (hpi) [18]. TauSTED super-resolution imaging of the sample revealed a close association between EmeraldGFP-Rab6a (EmGFP-Rab6a) and the Golgin97, validating that our EmGFP-Rab6a construct does localize to and serves as a marker of the *trans*-Golgi (Figure 1A).

**FIG 1.**
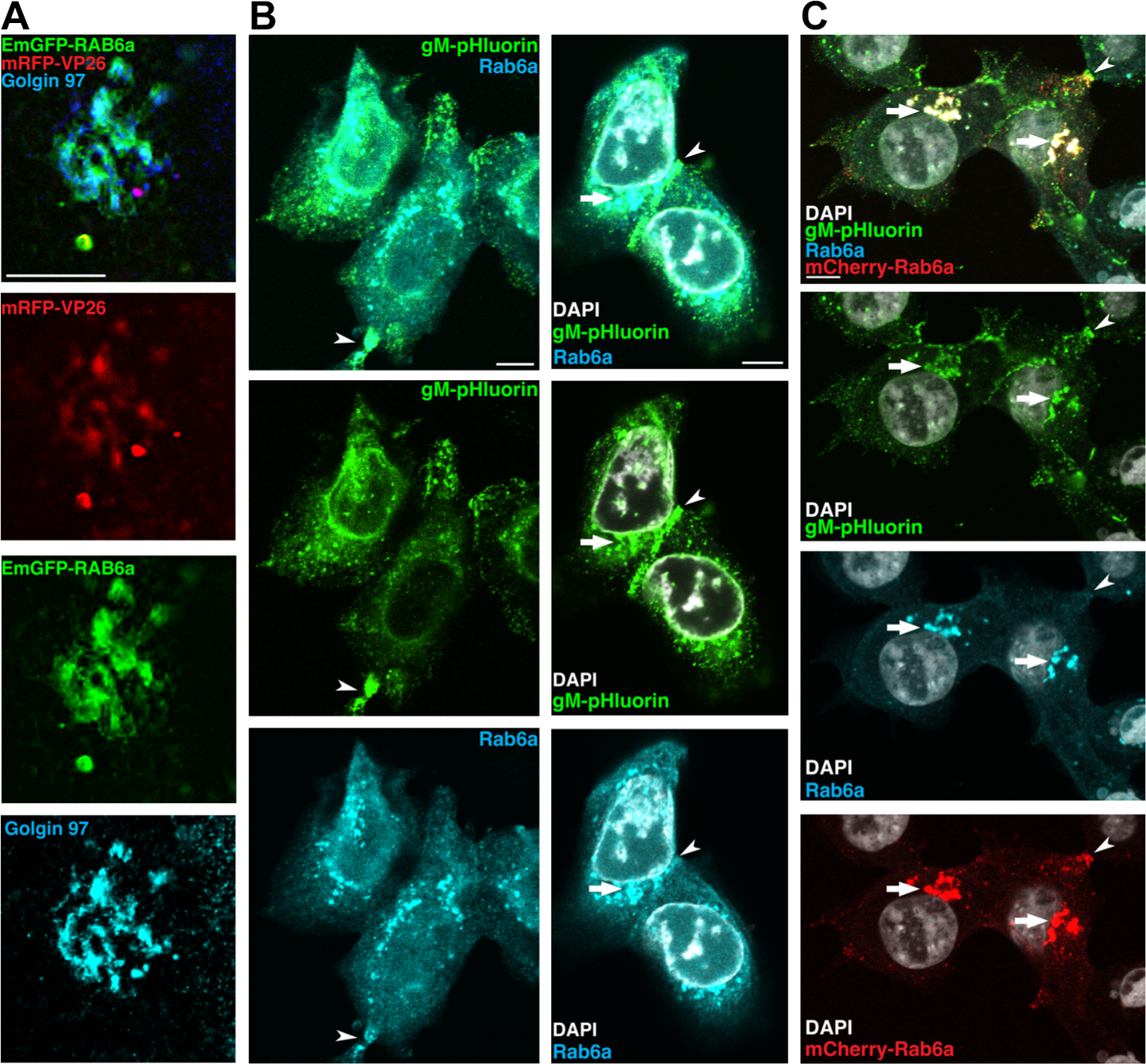
**A**. TauSTED superresolution imaging of HSV-1 mRFP-VP26 colocalizes with *trans*-Golgi (Golgin97) in Vero cells. Samples were infected with HSV-1 amplicon vector expressing EmGFP-Rab6a and HSV-1 OK14 helper virus. Samples were fixed and stained at 6hpi for Golgin97. Rab6a and Golgin97 colocalize, and HSV-1 capsid colocalizes with Golgin97 in the *trans*-Golgi. Scale bar represents 5*µ*m. **B.** Confocal microscopy of fixed and immunostained cells detecting endogenous Rab6a, with HSV-1 infection, in Vero cells at 7 hpi. Scale bars represents 10*µ*m. Endogenous Rab6a (blue) and HSV-1 gM-pHluorin protein (green) colocalizes to the juxtanuclear space and cell periphery in Vero cells that are not transduced to express exogenous Rab6a. **C.** Endogenous Rab6a (blue), exogenous mCherry-Rab6a (red), and HSV-1 gM-pHluorin (green) colocalize in Vero cells transduced with an adenovirus vector to express exogenous mCherry-Rab6a.

### Endogenous Rab6a Colocalizes with Exogenous Rab6a

To further explore the Rab6a association with the TGN and HSV-1, we performed a series of confocal microscopy experiments. In cells that were infected with HSV-1, but not transduced to express exogenous Rab6a, endogenous Rab6a was found in the juxtanuclear Golgi area, and with some accumulation at the cell periphery and cell-cell junctions (Figure 1B). Exogenous mCherry-Rab6a expressed from a non-replicating adenovirus vector also colocalized with endogenous Rab6a (Figure 1C). These results show that the colocalization and peripheral accumulations of Rab6a described below are not due to viral vector transduction and overexpression of fluorescent protein-tagged variants of Rab6a.

### Rab6a Colocalizes with HSV-1 Envelope and Capsid Proteins in the Golgi Region

Previously, we constructed a recombinant virus, HSV-1 IH01 expressing a pH-sensitive variant of GFP fused to the extravirion loop of glycoprotein M (gM-pHluorin) [19]. In fixed and immunostained cells, cellular pH equilibrates to that of the extracellular buffer, so gM-pHluorin exhibits consistent green fluorescence. At 6 hpi in Vero cells infected with HSV-1 IH01, gM-pHluorin appears in the juxtanuclear Golgi region, and colocalizes with endogenous Rab6a (Figure 1B) and exogenous mCherry-Rab6a (Figure 1C). We also observed clustering of gM-pHluorin at the cell periphery, similarly to the clustering of capsids we previously described [19] (Figure 1B-C). HSV-1 capsids, labeled with an mRFP-VP26 capsid protein, colocalize with gM-pHluorin and Rab6a in the juxtanuclear Golgi region [18], (Figure 1, Figure 2C), which is likely the site of secondary envelopment.

**FIG 2.**
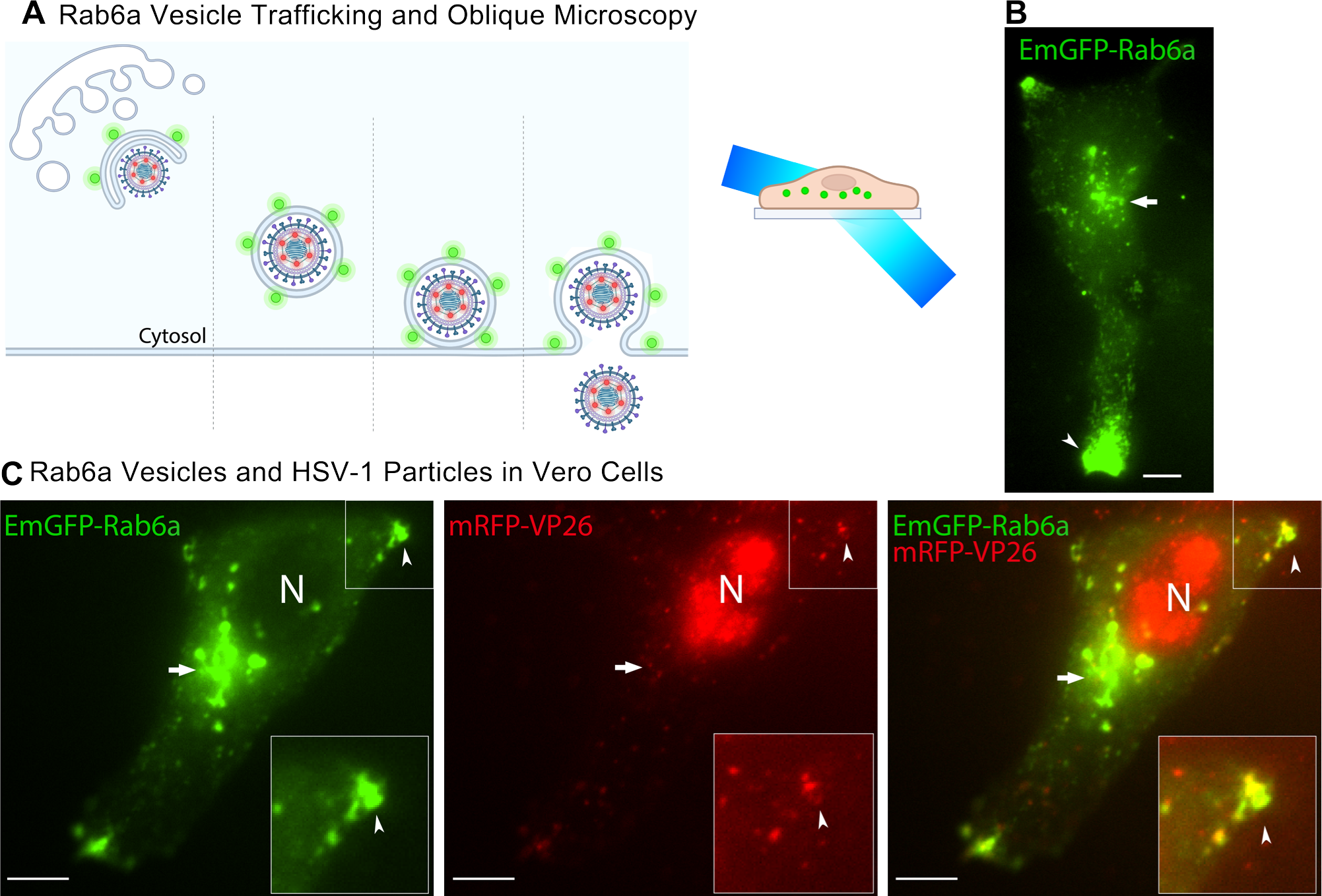
**A.** Schematic of viral egress and fluorescent reporters. Secretory vesicles marked with EmGFP-Rab6a (green) transport virus particles marked with mRFP-VP26 (red) to the plasma membrane. Oblique microscopy projects a light sheet that excites fluorescent molecules within a thin section of the cell volume. **B.** Vero cells transduced with an HSV-1 amplicon vector expressing EmGFP-Rab6a. Rab6a localizes to the juxtanuclear Golgi region (arrow), and accumulates forming clusters at the cell periphery (arrowhead) at 6 hpi. **C.** EmGFP-Rab6a (green) and HSV-1 capsids (red) colocalize in clusters at the cell periphery (arrow) in and Rab6a localizes near the Golgi (arrowhead) in Vero cells 6 hpi. Scale bars represent 10*µ*m.

### Rab6a Secretory Vesicles Accumulate at Peripheral Egress Sites

Previously, we and others have shown how HSV-1 particles exit from infected cells at preferential sites at the cell periphery [20, 21, 22, 23, 19]. Specifically, we showed that HSV-1 virus particles preferentially traffic to the egress site, gradually forming large accumulations of virus particles over time at these sites, and then complete exocytosis as individual virus particles at the plasma membrane [19]. To assess the intracellular localization and trafficking of HSV-1 and Rab6a, we transduced Vero cells with the HSV-1 EmGFP-Rab6a amplicon vector and imaged the cells with oblique microscopy. Sometimes called HiLo or “dirty TIRF”, oblique microscopy is similar to TIRF microscopy, except that the excitation laser is projected into a light sheet that selectively illuminates a thin section of the cell (Figure 2A). Using this method, we were able to project the light sheet to different depths of the cell by varying the incident angle of the excitation laser. By projecting the light sheet deeper into the cell with oblique microscopy, we observed that EmGFP-Rab6a strongly localizes to juxtanuclear organelles 6hpi in live cells that are singly infected with the EmGFFP-Rab6a amplicon and do not have detectable mRFP-VP26 capsid protein expression (Figure 2B), consistent results of the superresolution and confocal microscopy we performed (Figure 1). Rab6a also accumulates in clusters at the cell periphery in these uninfected cells, so peripheral Rab6a accumulation sites are not a result of productive HSV-1 infection (Figure 2B).

In cells that were doubly infected with both the EmGFP-Rab6a amplicon and HSV-1 OK14, large amounts of mRFP-VP26 capsids accumulate in the nucleus, a few particles colocalize with EmGFP-Rab6a in the juxtanuclear Golgi region, and particles begin to appear at sites of EmGFP-Rab6a clustering at the cell periphery by 6 hpi (Figure 2C). Importantly, virus particles and Rab6a vesicles appear to form accumulations in similar locations in singly and coinfected cells, indicating HSV-1 particles and Rab6a share similar trafficking routes and intracellular localization mechanisms.

### Colocalization of RabGTPases and HSV-1 in Human Fibroblast MRC5 Cells

In the interest of investigating the role of Rab6a in HSV-1 egress in a more biologically relevant cell type, we infected human MRC5 fibroblasts with the EmGFP-Rab6a amplicon and HSV-1 OK14 [24]. At 6 hpi, we imaged the cell with oblique microscopy. In MRC5 cells that were positive for only expression of EmGFP-Rab6a, whose nuclei did not have detectable mRFP-VP26 capsid protein to indicate HSV-1 infection, Rab6a accumulates at the cell periphery and in the juxtanuclear region (Figure 3A). As these cells are fibroblasts, they exhibit a longer phenotype than Vero cells, and the accumulation of Rab6a tends to occur at the tips of these cells (Figure 3A-C). In coinfected cells, we found that HSV-1 capsids colocalize in these peripheral tips with Rab6a, and capsids in the nucleus indicate robust HSV-1 infection (Figure 3B). We also transduced MRC5 cells with an adenovirus vector so that they expressed exogenous mCherry-Rab6a or mCherry-Rab27a. At 6hpi, we imaged the transduced cells with TIRF microscopy. In the absence of HSV-1 infection, Rab6a accumulates in the cell periphery while Rab27a does not (Figure 3C-D).

**FIG 3.**
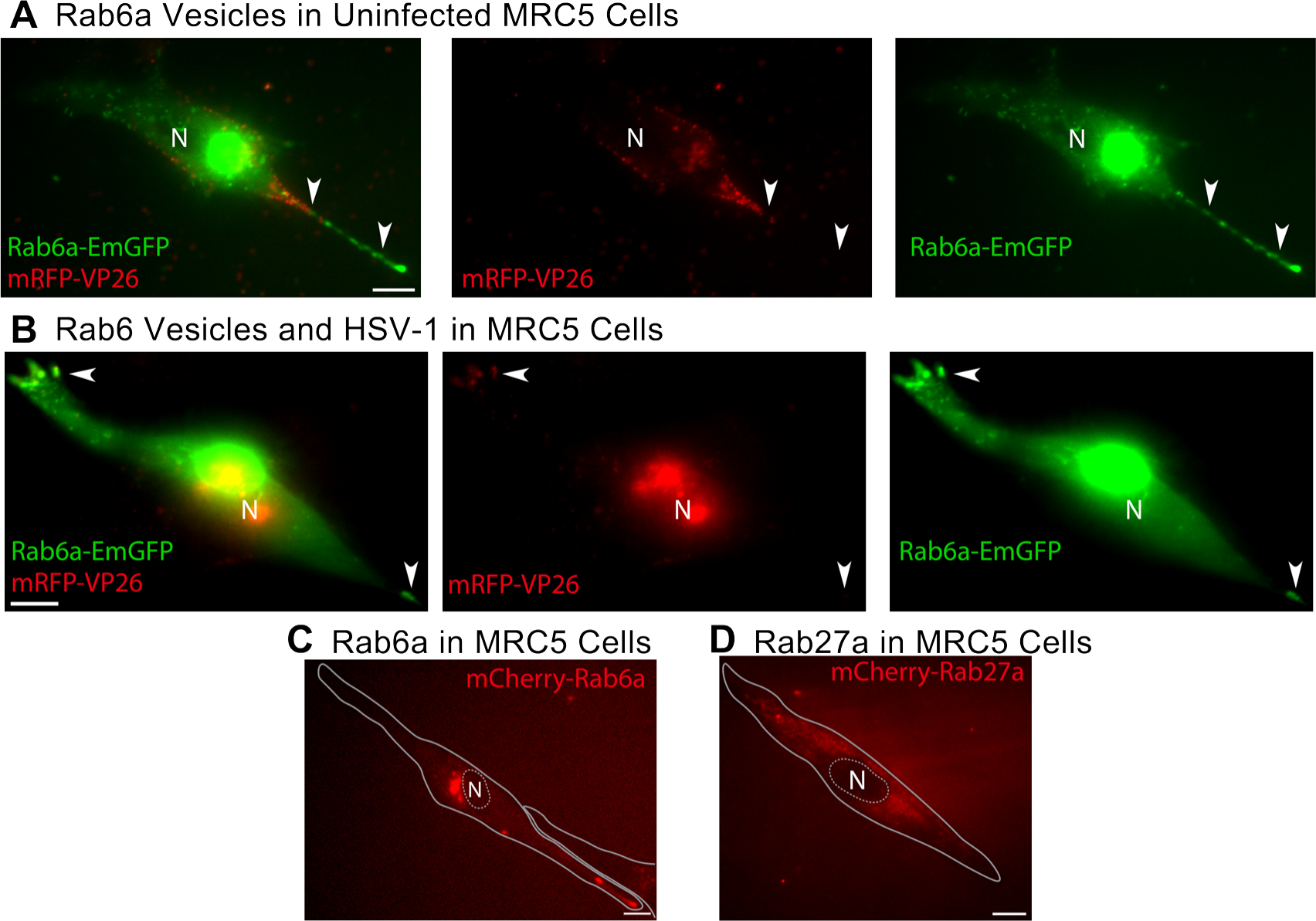
Oblique and TIRF microscopy of human lung fibroblast MRC5 cells expressing fluorescent Rab6a and coinfected with HSV-1 at 6hpi. Scale bar = 10*µ*m. **A.** Oblique microscopy of representative MRC5 cell transduced with EmGFP-Rab6a amplicon vector showing accumulations of EmGFP-Rab6a (green) in the juxtanuclear region and in the cell periphery (arrowheads). **B.** Oblique microscopy showing colocalization of EmGFP-Rab6a (green) and HSV-1 mRFP-VP26 capsid protein (red) in the cell periphery (arrowheads). **C.** TIRF microscopy of the accumulation of mCherry-Rab6a in the juxtanuclear region and cell periphery in cells transduced with an adenovirus vector. **D.** TIRF microscopy showing that Rab27a, expressed from an adenovirus vector, does not accumulate in the juxtanuclear region or in the cell periphery.

### HSV-1 Particles and Rab6a Vesicles Cotraffic from the Golgi Region to the Plasma Membrane

After exiting the nucleus, HSV-1 capsids transport to the cellular membranes where secondary envelopment occurs. While the organelles that contribute to HSV-1 secondary envelopment are not entirely clear, viral membrane proteins do traffic through the TGN, colocalize with viral tegument proteins and capsids, and therefore the TGN or TGN-derived secretory organelles are likely where secondary envelopment occurs [25, 26, 27, 28, 29].

To assess whether virus particles cotraffic with Rab6a, we coinfected Vero cells with HSV-1 OK14 and the EmGFP-Rab6a amplicon, and imaged by live-cell TIRF microscopy so that the cytoplasm near the plasma membrane was visible, but structures deeper in the cell, such as the nucleus, were largely excluded. Again, capsids were found in the juxtanuclear Golgi region, including colocalizing with discrete EmGFP-Rab6a puncta (Figure 4A). Over the course of several minutes, we observed EmGFP-Rab6a vesicles and mRFP-VP26 capsids traffic together towards the cell periphery, as shown using maximum difference projections and kymographs (Figure 4B-C) of the same infected cell shown in Figure 3A. Maximum difference projections highlight areas where fluorescence increases rapidly, while suppressing background fluorescence from structures that are not moving or changing over time. Kymographs show particle movement over distance on the X-axis and movement over time on the Y-axis. Two particles, marked as 1 and 2 in Figure 4B, that are positive for both EmGFP-Rab6a and mRFP-VP25 move together on their respective paths. The kymographs in Figure 4C show the spatial-temporal aspects of these two particles, revealing that the transport of HSV-1 particles in Rab6a secretory vesicles is a dynamic process across many micrometers from the interior of the cell to its periphery.

**FIG 4.**
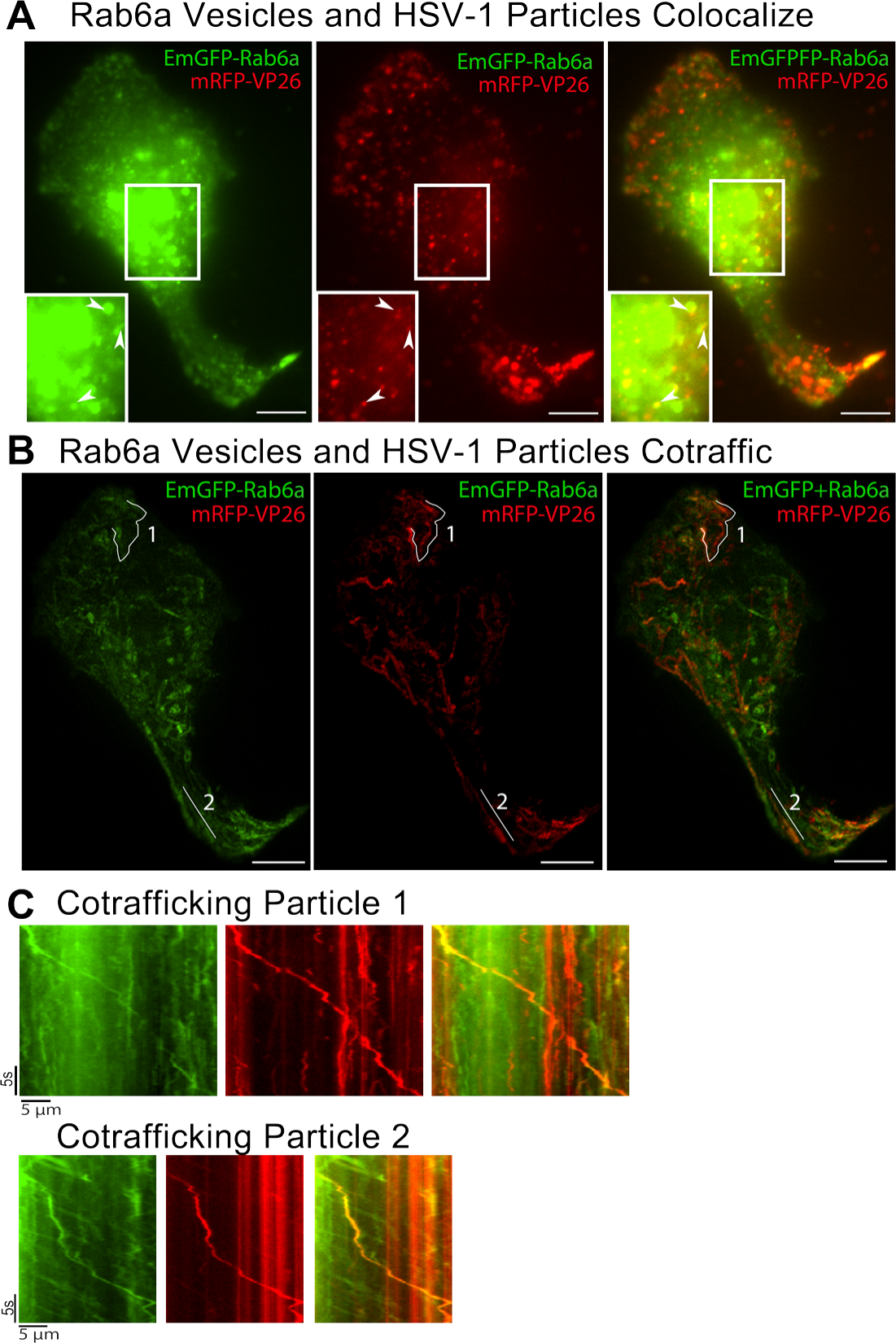
HSV-1 particles colocalize and cotraffic with Rab6a towards the cell periphery. Vero cells transduced with an HSV-1 amplicon vector expressing EmGFP-Rab6a, coinfected with HSV-1 OK14, and imaged by live-cell oblique microscopy at 6 hpi. Scale bars = 10*µ*m. **A.** HSV-1 capsids and EmGFP-Rab6a colocalize in distinct puncta in the Golgi region **B.** Rab6a and HSV-1 capsids cotraffic towards the cell periphery. **C.** Kymographs showing movement of Rab6a and HSV-1 capsid cotrafficking. Kymographs show movement of Particles 1 and 2 noted in panel B.

### HSV-1 Particles Undergo Exocytosis from Rab6a Vesicles

Since HSV-1 particles cotraffic with Rab6a to the plasma membrane, we next wanted to determine if these intracellular trafficking events culminate in virus particle exocytosis from these Rab6a vesicles. Previously, we showed that PRV particles undergo exocytosis from Rab6a vesicles in non-neuronal cells [11, 12] and from the cell body of primary neurons [30]. To visualize HSV-1 exocytosis, we infected Vero cells with HSV-1 IH01, which expresses gM-pHluorin. We previously showed that gM-pHluorin is incorporated into virus particles, and exhibits pH-dependent green fluorescence [19]. In live cells, the lumen of secretory vesicles is acidic (pH of 5.2-5.7) [31, 32], which quenches the green fluorescence of gM-pHluorin (Figure 5A). At the moment of exocytosis, the pHluorin moiety is exposed to the extracellular pH and its fluorescence is dequenched [31, 11, 12, 19], allowing us to identify virus particle exocytosis events using TIRF microscopy (Figure 5A).

**FIG 5.**
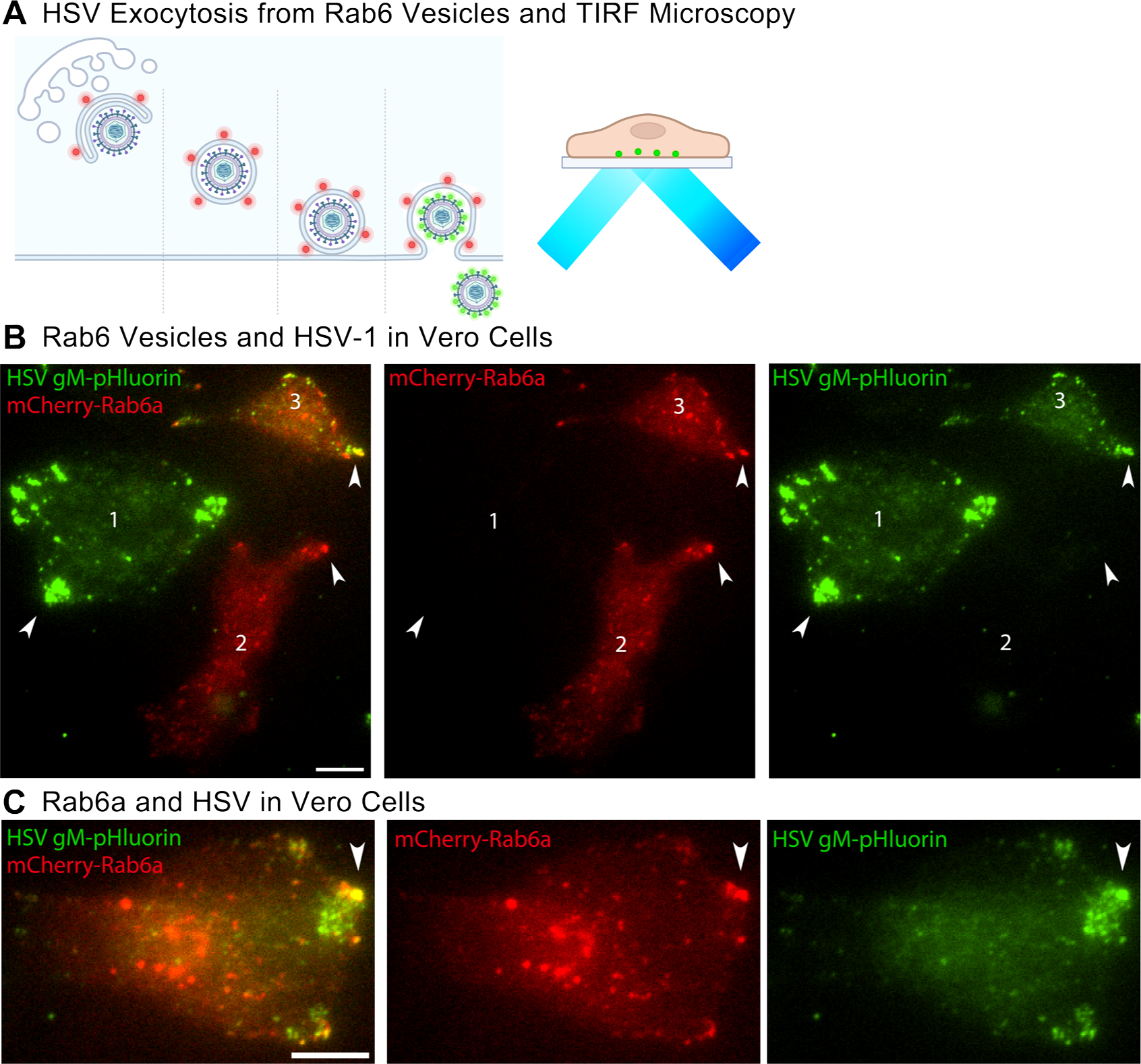
TIRF microscopy at 6 hpi of Vero cells coinfected with HSV-1 IH01 expressing gM-pHluorin (green) and an adenovirus vector expressing mCherry-Rab6a (red). Scale bars = 10*µ*m. **A.** Schematic of mCherry-Rab6a secretory vesicles transporting virus particles to the plasma membrane. gM-pHluorin becomes fluorescent with the pH change at the moment of exocytosis. TIRF microscopy excites fluorescent molecules at the plasma membrane. **B.** HSV-1 gM accumulates in the cell periphery in cells that are not transduced (1) and are transduced (3). Rab6a accumulates in the cell periphery in cells that are not transduced (2) and are transduced (3). **C.** HSV-1 particles undergo exocytosis from Rab6a secretory vesicles at the cell periphery, and HSV-1 and Rab6a accumulate at these locations in the infected cell.

Vero cells were transduced with a non-replicating adenovirus vector expressing mCherry-Rab6a, coinfected with HSV-1 IH01, and imaged by live-cell TIRF microscopy beginning at 6 hpi. HSV-1 particles containing gM-pHluorin accumulated at peripheral clusters in cells that were infected only with HSV-1 and were not positive for mCherry-Rab6a (Figure 5B, Cell 1). Cells that were singly transduced, with expression of mCherry-Rab6a but no detectable HSV-1 gM-pHluorin, showed accumulations of Rab6a in peripheral clusters (Figure 5B, Cell 2). In cells that were positive for both HSV-1 gM-pHluorin and mCherry-Rab6a, both proteins accumulated in the same locations in the cell periphery (Figure 5, Cell 3, and Figure 5C).

After exocytosis, virus particles remained attached to the cell surface and largely immobile, resulting in the accumulation of large clusters of particles. Clusters of mCherry-Rab6a most likely represent secretory vesicles that have not yet fused to the plasma membrane, and clusters of dequenched, fluorescent gM-pHluorin puncta represent a combination of virions and L-particles. Because we also observed similar clustering of mRFP-VP26 capsids (Figures 2 and 4), and clusters of colocalized gM-pHluorin and mRFP-VP26 capsids [19], a subset of these clustered particles represent complete, mature virions.

### Quantification of Individual Exocytosis Events from Rab6a Vesicles

In Vero cells, identifying exocytosis of virus particles from individual Rab6a vesicles was difficult against the high background fluorescence from these large clusters of previously released particles - individual vesicles were no longer distinguishable in a large cluster. To overcome this problem, we transduced and infected PK15 (porcine kidney epithelial) cells, which were previously used to investigate egress of PRV [11, 12]. We have shown that HSV-1 productively infects PK15 cells, but unlike in Vero cells, HSV-1 and PRV exocytosis events are uniformly distributed across the adherent cell surface, with much less accumulation of large clusters [19].

Over the course of several minutes, we observed individual virus particles undergoing exocytosis from mCherry-Rab6a vesicles, which were easily distinguishable without the formation of large clusters. A representative exocytosis event is shown using maximum difference projection and a kymograph (Figure 6A). Here, maximum difference projections highlight areas where fluorescence increases rapidly due to gM-pHluorin dequenching, and kymographs show the change in particle fluorescence over time on the Y-axis.

**FIG 6.**
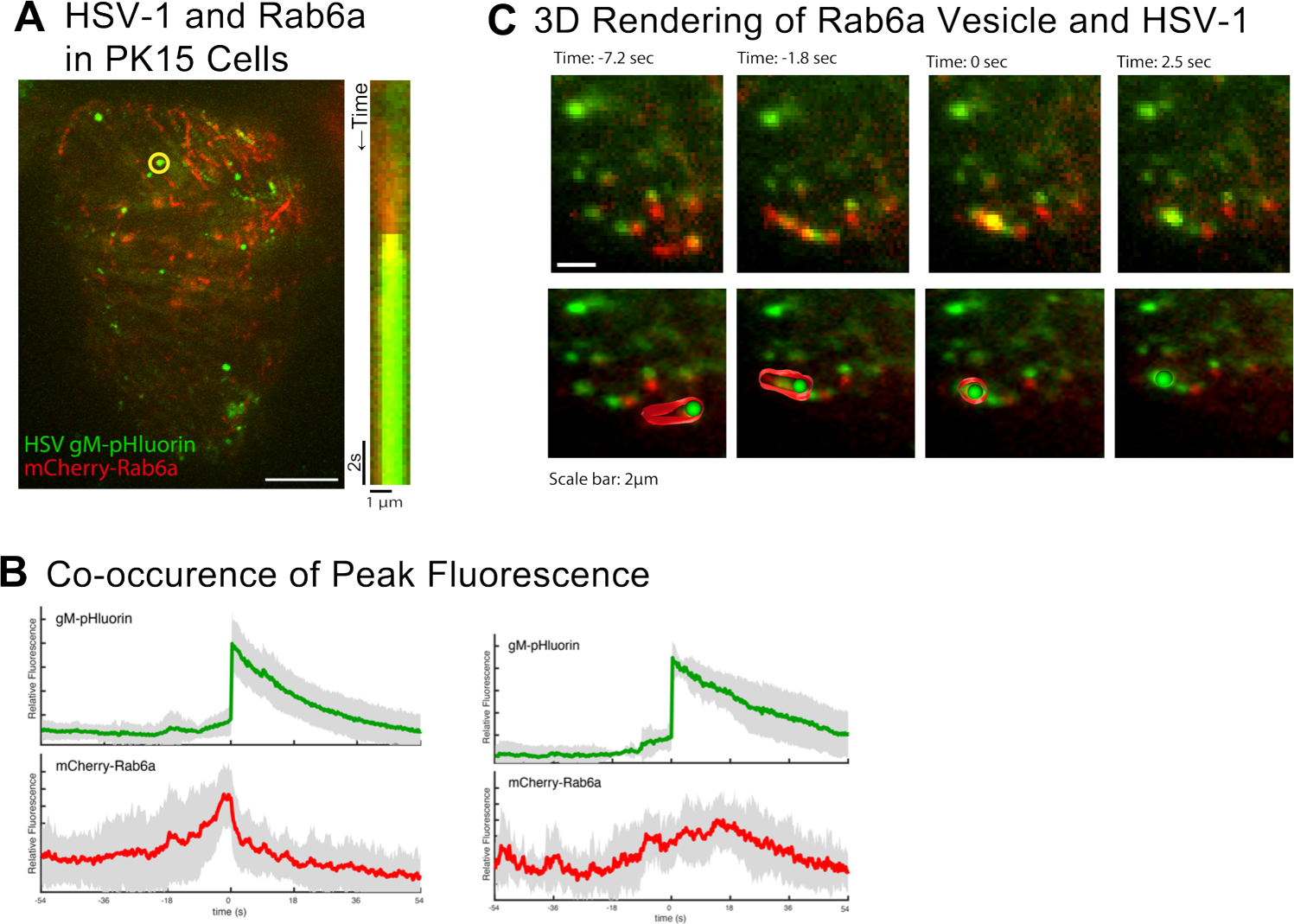
TIRF microscopy at 6 hpi of PK15 cells transduced with mCherry-Rab6a adenovirus vector and infected with HSV-1 IH01. **A.** Maximum difference projection of coinfected cell over 3:38sec of imaging. Kymograph of particle (yellow circle) movement and fluorescence over time showing HSV-1 exocytosis from a Rab6a vesicle. Scale bar = 10*µ*m. **B.** Plot of mean fluorescence over time at exocytosis event (time = 0). mCherry-Rab6a fluorescence (red) peaks before HSV-1 gM-pHluorin (green) as the secretory vesicle arrives and the plasma membrane and the virus particle is released to the extracellular space. Shading is standard deviation (n=44). **D**. Rab6a vesicle transporting a virus particle to the plasma membrane for viral exocytosis, rounded up for viral exocytosis, and dissipated after viral exocytosis. Top row shows still frame images at the indicated time points before, at, and after the exocytosis event (Time=0 sec). Bottom row shows 3D model, tilted at *∼*36°on the X-axis, of the Rab6a vesicle and virus particle compiled in Imaris for the same time points. Scale bar represents 2*µ*m.

To quantify gM-pHluorin and mCherry-Rab6a fluorescence over many individual exocytosis events, we calculated the average relative fluorescence intensity over time. Individual exocytosis events were measured, and time series data were aligned to a common time=0 based on the peak of gM-pHluorin fluorescence intensity. Average green fluorescence exhibits a sharp increase, representing rapid dequenching of gM-pHluorin upon exocytosis, followed by a slow exponential decay due to photobleaching. For a majority of exocytosis events (n=27/44, 61%), red fluorescence gradually increased prior to exocytosis, representing the arrival of a mCherry-Rab6a positive vesicle to the site of exocytosis. Immediately after exocytosis, the red signal rapidly decays, representing a combination of mCherry-Rab6a molecules diffusing away from the site of exocytosis and photobleaching (Figure 6B).

However, some exocytosis events (n=17/44, 39%) did not appear to be associated with a strong mCherry-Rab6a signal (Figure 6C). This may be a result of low mCherry-Rab6a expression, high fluorescence background, or a minority of HSV-1 particles may use alternative secretory mechanisms.

### Secretory Rab27a and Endocytic Rab5a and Are Not Associated with HSV-1 Egress

As negative controls, we also tested Rab27a and Rab5a. Many enveloped viruses, including HSV-1 [33], use cellular ESCRT machinery for virion budding or membrane scission. In uninfected cells, ESCRT produces intralumenal vesicles in late endosomes/multivesicular bodies. Rab27a regulates the exocytosis of endosomal organelles, including multivesicular bodies, to produce extracellular exosomes. Rab27 has been implicated in HSV-1 secondary envelopment, specifically in an oligodendrocyte cell line where Rab27 appears to co-localize with the trans-Golgi [34]. Rab5 is present on early endosomes, and has also been implicated in HSV-1 secondary envelopment [13, 14].

To investigate the role of these Rab GTPases, we transduced Vero cells with adenovirus vectors expressing mCherry-Rab27a or mCherry-Rab5a, then infected with HSV-1 expressing gM-pHluorin, and imaged by TIRF microscopy beginning at 6 hpi. As shown in Figure 7A, HSV-1 particles undergo exocytosis from mCherry-Rab27a transduced cells (circles), and mCherry-Rab27a localizes to discrete clusters (boxes). However, mCherry-Rab27a does colocalize with gM-pHluorin (Figure 7A), and does not appear to be associated with viral exocytosis (Figure 7C). Similarly, mCherry-Rab5a localizes to distinct puncta scattered throughout the cell, but does not appear to strongly colocalize with gM-pHluorin (Figure 7B), and does not appear to be associated with viral exocytosis (Figure 7C). These results show that we do not detect spurious colocalization/egress with any Rab GTPase due to exogenous expression. While Rab27 may play important roles in specialized cell types, and Rab5 may be important for viral glycoprotein trafficking upstream of HSV-1 egress, these Rabs do not appear to be associated with viral exocytosis.

**FIG 7.**
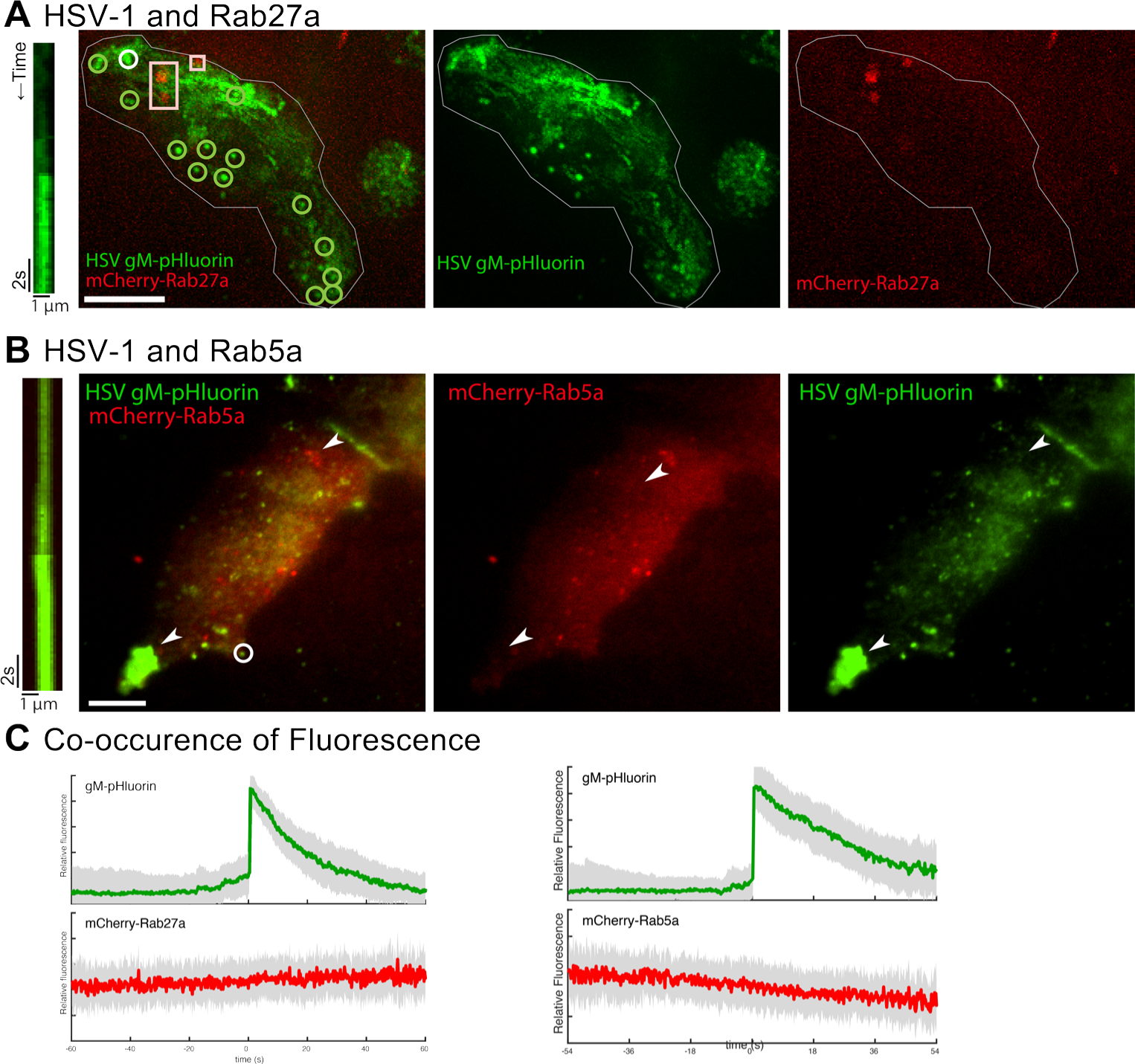
TIRF microscopy at 6 hpi of Vero cells transduced with mCherry-Rab27a adenovirus vector (red) and infected with HSV-1 IH01 expressing gM-pHluorin (green). Scale bars = 10*µ*m. **A.** Representative cell expressing mCherry-Rab27a and infected with HSV-1 IH01. Individual virus particle exocytosis events are marked with green circles, and Rab27a vesicles are marked with red boxes. The viral exocytosis event illustrated in the kymograph is marked with a white circle. Virus particle exocytosis events occur in preferential sites in the cell, and Rab27a is not found in these preferential exocytosis sites **B.** Green fluorescence peaks at the moment of exocytosis and virus particle release at the plasma membrane, whereas red fluorescence has no peak at that time. Shading is standard deviation. (n=55 exocytosis events from 10 cells). **C.**. HSV-1 does not undergo exocytosis from mCherry-Rab5a vesicles in Vero cells. Scale bar = 10*µ*m. HSV-1 gM-pHluorin accumulates in hotspots and cell junctions. Rab5a does not accumulate with HSV-1 progeny or envelope proteins. **D.**. Plot of mean HSV-1 gM-pHluorin and mCherry-Rab27a or mCherry-Rab5a, respectively, fluorescence aligned at the exocytosis event (time=0). gM-pHluorin fluorescence peaks at exocytosis. mCherry-Rab27a and mCherry-Rab5a have no corresponding peak with exocytosis. Shading is standard deviation (n=63 exocytosis events).

## DISCUSSION

Viral egress is a highly involved process: after assembly, virus particles must be sorted and delivered to the appropriate site of exocytosis, and must be released from the infected cell by exocytosis. It has been shown via knockdown and knockout studies that Rab6a is important for HSV-1 replication. We have also previously shown that Rab6a secretory vesicles are involved in egress of the related alpha herpesvirus, PRV [13, 29, 11, 12, 30]. By using live cell fluorescence microscopy, we found that Rab6a is associated with the final stages of HSV-1 replication. Rab6a and HSV-1 progeny particles colocalize in the region of the Golgi (Figures 1-2), where they cotraffic to the plasma membrane (Figure 4). At the plasma membrane, virus particles undergo exocytosis from Rab6a secretory vesicles and Rab6a diffuses from the exocytosis site while the virus particle remains (Figure 5-6).

However, Rab6a is likely not the sole Rab GTPase that plays a role in HSV-1 egress and exocytosis. PRV particles undergo exocytosis from secretory vesicles that are also labeled with Rab8a and Rab11a [11, 12], and the same might be the case for HSV-1. Rab6a is also not the only effector for microtubule transport and cargo trafficking, and it interacts with Rab8 and Rab11 [3, 35]. Since we found that some viral exocytosis events were not associated with detectable Rab6a, it may be that these viral egress events were associated with other Rab proteins.

Furthermore, alpha herpesviruses have evolved to carry out their replication cycles in a broad range of host cells, including neurons with highly specialized cell biology. Thus, these viruses have to overcome differences of host cell biology from one cell type to another. With HSV-1 specifically, it can be expected that there is some overlap between host factors involved with egress across cell types, as humans express several dozen different types of Rab GTPases [1]. Rab6a is expressed in neurons, along with a Rab6b isoform that is neuron-specific [36]. It is possible that both isoforms of Rab6 may contribute to HSV-1 transport and egress in neuronal cells.

In addition to HSV-1 egress from Rab6a vesicles, we also observed that HSV-1 particles and Rab6a accumulated in clusters together at the cell periphery. This clustering has been previously reported by others [20, 21, 22], including as far back as some of the earliest indirect immunofluorescence studies with HSV-1 [23]. In the cell biological literature, it has been shown that Rab6a coordinates the delivery of secretory vesicles near focal adhesions [5]. The coclustering of HSV-1 particles and Rab6a may be explained by this function of Rab6a targeting exocytosis to particular sites in the cell. Further studies will be needed to determine what connections may exist between Rab6, other Rab GTPases that may function combinatorially, and their effector molecules that govern trafficking and exocytosis.

## MATERIALS AND METHODS

### Cell lines and cell culture

Vero (ATCC, CCL-81), human lung fibroblast MRC5 (ATTC, CCL-171),and PK15 (ATCC, CCL-33) cells were obtained from ATCC, and maintained in DMEM medium (Cytiva) supplemented with 10% fetal bovine serum (Omega and Peak Scientific) and 1% penicillin-streptomycin (Gibco), in a 5% *CO*_2_ incubator at 37°C. HEK293 cells were a kind gift from Masmudur Rahman (Arizona State University).

### HSV-1 Viruses

HSV-1 IH01 was constructed and propagated as previously described [19]. HSV-1 OK14 was obtained from Lynn Enquist (Princeton University) [17] from whole-genome sequenced archival stocks.

### HSV-1 Amplicon Vector Construction and Propagation

The amplicon vector plasmid pCPD-HSV-N-EmGFP-DEST was constructed by the DNASU Plasmid Repository (Biodesign Institute, Arizona State University), as follows: The plasmid HSV-DYN-hM4Di was a gift from John Neumaier (Addgene plasmid # 53327) [37]. Unnecessary promoter and transgene sequences were removed by digestion with HindIII and religating, to produce pCPD-HSV. This plasmid contains a HSV-1 packaging signal and OriS origin of replication. The plasmid pcDNA6.2/N-EmGFP-DEST obtained from Invitrogen (ThermoFisher). The CMV promoter, Emerald GFP coding region, Gateway recombination cassette (attR1, CmR selection marker, ccdB counterselection marker, attR2), and HSV-1 TK polyadenylation signal were PCR amplified and ligated into the HindIII site on pCPD-HSV, to produce pCPD-HSV-N-EmGFP-DEST. The human Rab6a coding sequence (DNASU Plasmid Repository, #HsCD00296778, NCBI Nucleotide reference BC096818) was inserted by Gateway recombination (Invitrogen) to make an in-frame EmGFP-Rab6a fusion.

To propagate the EmGFP-Rab6a amplicon vector, 3.5 × 10^5^ HEK 293A cells were seeded into each well of a 6-well plate (Celltreat), incubated overnight, and then transfected with 3*µ*g of amplicon plasmid using Lipofectamine 2000 (Invitrogen). 24 hours after transfection, cells were infected with 10^5^ infectious units of HSV-1 OK14. Cells were incubated for another 24 hours, and then cells and supernatants were harvested. The amplicon stock was passaged at high MOI three times on Vero cells, cells and supernatants were harvested, and stored at −80°C.

### Adenovirus Vectors

Non-replicating E1/E3-deleted adenovirus vectors expressing mCherry-Rab6a or mCherry-Rab5a were previously described [11, 12], and propagated on HEK 293A cells.

### Live Cell Oblique and TIRF Microscopy

Cultures were prepared for TIRF and oblique microscopy by seeding cells at a low density on glass-bottom 35mm dishes (Celltreat, Ibidi, or MatTek), and incubated overnight before viral transduction and infection. Cells were infected with HSV-1, amplicon vectors, and/or adenovirus vectors, and imaged beginning at 6 hpi. The HSV-1, amplicon vector, and adenovirus vector inoculum amounts were determined empirically to synchronously infect/transduce a majority of cells as observed by fluorescence microscopy.

Fluorescence microscopy was performed using a Nikon Eclipse Ti2-E inverted microscope in the Biodesign Imaging Core facility (Arizona State University, Tempe, AZ). This microscope is equipped with TIRF and widefield illuminators, a Photometrics Prime95B sCMOS camera, a 60X high-NA TIRF objective, and objective and stage warmers for 37°C live-cell microscopy. 488nm and 561nm lasers were used to excite green and red fluorescent proteins, respectively.

### Immunofluorescence and Confocal Microscopy

Vero cells were seeded into 8-well Ibidi dishes and cultured overnight. Cells were infected with HSV-1 IH01, mCherry-Rab6a adenovirus vector, or cotransduced/infected, as described above. At 7 hpi, cells were fixed for 10 minutes with 4% freshly-prepared paraformaldehyde in PBS, permeabilized with 0.5% Triton X-100 for 90 seconds, then fixed again for 2 minutes. After fixation, samples were rinsed three times with PBS, then blocked with 5% goat serum in PBS for 30 minutes at room temperature. Primary rabbit polyclonal antibody against Rab6a (Abcam ab95954) was used at a 1:200 dilution in antibody diluent (0.05% Tween 20, 1% goat serum, in PBS). Cells were incubated in primary antibody overnight in the dark at 4°C, then rinsed 3 times in wash buffer (0.05% Tween 20, in PBS). Secondary antibody, anti-rabbit AlexaFluor 633 (ThermoFisher A21071) was diluted 1:1000 in antibody diluent and incubated on samples for 2 hr at room temperature with gentle rocking. Cells well rinsed 3 times with wash buffer, and once with PBS. Nuclei were labeled with 0.1 *µ*g/mL DAPI in PBS. Samples were imaged on a Nikon AX R laser scanning confocal microscope in the Biodesign Imaging Core facility (Arizona State University, Tempe, AZ) using a 60X 1.42 NA objective. The DAPI channel was excited at 405nm, pHluorin at 488nm, mCherry at 568nm, and AlexaFluor 633 at 640nm. Emissions for these channels were collected in the blue, green, red, and far red spectral ranges respectively.

### Image Analysis

Image analysis to identify exocytosis events and HSV colocalization with Rab6a vesicles was conducted in Fiji software [38]. Fluorescence microscopy images were prepared for publication using Adjust Brightness/Contrast, Reslice (to produce kymographs), and Plot Z-axis Profile (to measure fluorescence over time) functions in Fiji. Maximum difference projections were calculated as previously described [11], using the Duplicate, Stacks->Tools, Math->Subtract, and Z Project functions in Fiji. Maximum difference projection shows where fluorescence intensity increases most rapidly, which emphasizes exocytosis events and particle movement, and deemphasizes static features that do not change during the course of imaging. Plots of average fluorescence intensity during exocytosis events were calculated using Matlab (Mathworks). Image segmentation and 3D rendering of HSV-1 virus particle undergoing exocytosis from Rab6a vesicle was done in Imaris (Oxford Instruments) by constructing spots and surfaces for the objects at respective image slices. Access to the Imaris software was kindly provided by Drs. M. Foster Olive and Jessica Verpeut (Dept. of Psychology, Arizona State University).

## ACKNOWLEDGMENTS

Thank you to Joli Bastin for her work on producing the recombinant HSV IH01 and IH02 strains carrying gM-pHluorin.

## DATA AVAILABILITY STATEMENT

All data and regeants available upon request.

## References

[1] Stenmark H. 2009. Rab GTPases as coordinators of vesicle traffic. Nature Reviews Molecular Cell Biology 10:513–525. doi:10.1038/nrm2728.

[2] Goud B, Zahraoui A, Tavitian A, Saraste J. 1990. Small GTP-binding protein associated with Golgi cisternae. Nature 345:553–556. doi:10.1038/345553a0.

[3] Grigoriev I, Splinter D, Keijzer N, Wulf PS, Demmers J, Ohtsuka T, Modesti M, Maly IV, Grosveld F, Hoogenraad CC, Akhmanova A. 2007. Rab6 Regulates Transport and Targeting of Exocytotic Carriers. Developmental Cell 13:305–314. doi: 10.1016/j.devcel.2007.06.010.

[4] Jordens I, Marsman M, Kuijl C, Neefjes J. 2005. Rab proteins, connecting transport and vesicle fusion. Traffic 6:1070–1077. doi:10.1111/j.1600-0854.2005.00336.x.

[5] Fourriere L, Kasri A, Gareil N, Bardin S, Bousquet H, Pereira D, Perez F, Goud B, Boncompain G, Miserey-Lenkei S. 2019. RAB6 and microtubules restrict protein secretion to focal adhesions. Journal of Cell Biology 218 (7):2215–2231. doi: 10.1083/jcb.201805002.

[6] Bär S, Daeffler L, Rommelaere J, Nüesch JPF. 2008. Vesicular Egress of Non-Enveloped Lytic Parvoviruses Depends on Gelsolin Functioning. PLOS Pathogens 4:e1000126. doi:10.1371/journal.ppat.1000126.

[7] Brass AL, Dykxhoorn DM, Benita Y, Yan N, Engelman A, Xavier RJ, Lieberman J, Elledge SJ. 2008. Identification of host proteins required for HIV infection through a functional genomic screen. Science 319:921–926. doi: 10.1126/science.1152725.

[8] Indran SV, Ballestas ME, Britt WJ. 2010. Bicaudal D1-Dependent Trafficking of Human Cytomegalovirus Tegument Protein pp150 in Virus-Infected Cells. Journal of Virology 84:3162–3177. doi:10.1128/jvi.01776-09.

[9] Indran SV, Britt WJ. 2011. A Role for the Small GTPase Rab6 in Assembly of Human Cytomegalovirus. Journal of Virology 85:5213–5219. doi:10.1128/jvi.02605-10.

[10] Girsch JH, Jackson W, Carpenter JE, Moninger TO, Jarosinski KW, Grose C. 2020. Exocytosis of Progeny Infectious Varicella-Zoster Virus Particles via a Mannose-6-Phosphate Receptor Pathway without Xenophagy following Secondary Envelopment. Journal of Virology 94. doi:10.1128/jvi.00800-20.

[11] Hogue IB, Bosse JB, Hu JR, Thiberge SY, Enquist LW. 2014. Cellular Mechanisms of Alpha Herpesvirus Egress: Live Cell Fluorescence Microscopy of Pseudorabies Virus Exocytosis. PLoS Pathogens 10:e1004535. doi:10.1371/journal.ppat.1004535.

[12] Hogue IB, Scherer J, Enquist LW. 2016. Exocytosis of Alphaherpesvirus Virions, Light Particles, and Glycoproteins Uses Constitutive Secretory Mechanisms. mBio 7:e00820–16. doi:10.1128/mbio.00820-16.

[13] Johns HL, Gonzalez-Lopez C, Sayers CL, Hollinshead M, Elliott G. 2014. Rab6 Dependent Post-Golgi Trafficking of HSV1 Envelope Proteins to Sites of Virus Envelopment. Traffic (Copenhagen, Denmark) 15:157–178. doi:10.1111/tra.12134.

[14] Hollinshead M, Johns HL, Sayers CL, Gonzalez-Lopez C, Smith GL, Elliott G. 2012. Endocytic tubules regulated by Rab GTPases 5 and 11 are used for envelopment of herpes simplex virus. The EMBO Journal 31:4204–4220. doi:10.1038/emboj.2012.262.

[15] Chavrier P, Goud B. 1999. The role of ARF and Rab GTPases in membrane transport. Current Opinion in Cell Biology 11 (4):466–475. doi:10.1016/s0955-0674(99)80067-2.

[16] Goud B, Liu S, Storrie B. 2018. Rab proteins as major determinants of the Golgi complex structure. Small GTPases 9 (1-2):66–75. doi:10.1080/21541248.2017.1384087.

[17] Song R, Koyuncu OO, Greco TM, Diner BA, Cristea IM, Enquist LW. 2016. Two Modes of the Axonal Interferon Response Limit Alphaherpesvirus Neuroinvasion. mBio 7:e02145–15. doi:10.1128/mbio.02145-15.

[18] Alzhanova D, Hruby DE. 2006. A trans-Golgi Network Resident Protein, golgin-97, Accumulates in Viral Factories and Incorporates into Virions during Poxvirus Infection. Journal of Virology 80 (23):11520–11527. doi:10.1128/jvi.00287-06.

[19] Bergeman MH, Hernandez MQ, Diefenderfer J, Drewes JA, Velarde K, Tierney WM, Hogue IB. 2023. Live-cell Fluorescence Microscopy of HSV-1 Cellular Egress by Exocytosis. bioRxiv p 2023.02.27.530373. doi:10.1101/2023.02.27.530373.

[20] Mingo RM, Han J, Newcomb WW, Brown JC. 2012. Replication of herpes simplex virus: egress of progeny virus at specialized cell membrane sites. Journal of virology 86:7084–97. doi:10.1128/jvi.00463-12.

[21] Johnson DC, Webb M, Wisner TW, Brunetti C. 2001. Herpes Simplex Virus gE/gI Sorts Nascent Virions to Epithelial Cell Junctions, Promoting Virus Spread. Journal of Virology 75:821–833. doi:10.1128/jvi.75.2.821-833.2001.

[22] McMillan TN, Johnson DC. 2001. Cytoplasmic Domain of Herpes Simplex Virus gE Causes Accumulation in the trans-Golgi Network, a Site of Virus Envelopment and Sorting of Virions to Cell Junctions. Journal of Virology 75:1928–1940. doi: 10.1128/jvi.75.4.1928-1940.2001.

[23] Norrild B, Virtanen I, Lehto VP, Pedersen B. 1983. Accumulation of herpes simplex virus type 1 glycoprotein D in adhesion areas of infected cells. Journal of General Virology 64:2499–2503. doi:10.1099/0022-1317-64-11-2499/cite/refworks.

[24] Jacobs JP, Jones CM,Baille JP. 1970. Characteristics of a Human Diploid Cell Designated MRC-5. Nature 227 (5254):168–170. doi:10.1038/227168a0.

[25] Turcotte S, Letellier J, Lippé R. 2005. Herpes Simplex Virus Type 1 Capsids Transit by the trans-Golgi Network, Where Viral Glycoproteins Accumulate Independently of Capsid Egress. Journal of Virology 79:8847–8860. doi:10.1128/jvi.79.14.8847-8860.2005.

[26] Sugimoto K, Uema M, Sagara H, Tanaka M, Sata T, Hashimoto Y, Kawaguchi Y. 2008. Simultaneous Tracking of Capsid, Tegument, and Envelope Protein Localization in Living Cells Infected with Triply Fluorescent Herpes Simplex Virus 1. Journal of Virology 82:5198–5211. doi:10.1128/jvi.02681-07.

[27] White S, Kawano H, Harata NC, Roller RJ. 2020. Herpes Simplex Virus Organizes Cytoplasmic Membranes To Form a Viral Assembly Center in Neuronal Cells. Journal of Virology 94 (19). doi:10.1128/jvi.00900-20.

[28] Miranda-Saksena M, Boadle RA, Aggarwal A, Tijono B, Rixon FJ, Diefenbach RJ, Cunningham AL. 2009. Herpes simplex virus utilizes the large secretory vesicle pathway for anterograde transport of tegument and envelope proteins and for viral exocytosis from growth cones of human fetal axons. Journal of virology 83:3187–99. doi: 10.1128/jvi.01579-08.

[29] Raza S, Alvisi G, Shahin F, Husain U, Rabbani M, Yaqub T, Anjum AA, Sheikh AA, Nawaz M, Ali MA. 2018. Role of Rab GTPases in HSV-1 infection: Molecular understanding of viral maturation and egress. Microbial Pathogenesis 118:146–153. doi: 10.1016/j.micpath.2018.03.028.

[30] Ambrosini AE, Deshmukh N, Berry MJ, Enquist LW, Hogue IB. 2019. Alpha Herpesvirus Egress and Spread from Neurons Uses Constitutive Secretory Mechanisms Independent of Neuronal Firing Activity. bioRxiv p 729830. doi: 10.1101/729830.

[31] Sankaranarayanan S, Angelis DD, Rothman JE, Ryan TA. 2000. The Use of pHluorins for Optical Measurements of Presynaptic Activity. Biophysical Journal 79:2199–2208. doi:10.1016/s0006-3495(00)76468-x.

[32] Paroutis P, Touret N, Grinstein S. 2004. The pH of the Secretory Pathway: Measurement, Determinants, and Regulation. Physiology 19:207–215. doi: 10.1152/physiol.00005.2004.

[33] Pawliczek T, Crump CM. 2009. Herpes Simplex Virus Type 1 Production Requires a Functional ESCRT-III Complex but Is Independent of TSG101 and ALIX Expression. Journal of Virology 83 (21):11254–11264. doi:10.1128/jvi.00574-09.

[34] Bello-Morales R, Crespillo AJ, Fraile-Ramos A, Tabarés E, Alcina A, López-Guerrero JA. 2012. Role of the small GTPase Rab27a during Herpes simplex virus infection of oligodendrocytic cells. BMC Microbiology 12:265. doi: 10.1186/1471-2180-12-265.

[35] Miserey-Lenkei S, Waharte F, Boulet A, Cuif MH, Tenza D, Marjou AE, Raposo G, Salamero J, Héliot L, Goud B, Monier S. 2007. Rab6-interacting Protein 1 Links Rab6 and Rab11 Function. Traffic 8:1385–1403. doi:10.1111/j.1600-0854.2007.00612.x.

[36] Opdam FJM, Echard A, Croes HJE, Hurk JAJMvd, Vorstenbosch RAvd, Ginsel LA, Goud B, Fransen JAM. 2000. The small GTPase Rab6B, a novel Rab6 subfamily member, is cell-type specifically expressed and localised to the Golgi apparatus. Journal of Cell Science 113:2725–2735. doi:10.1242/jcs.113.15.2725.

[37] Ferguson SM, Eskenazi D, Ishikawa M, Wanat MJ, Phillips PEM, Dong Y, Roth BL, Neumaier JF. 2010. Transient neuronal inhibition reveals opposing roles of indirect and direct pathways in sensitization. Nature Neuroscience 2010 14:1 14:22–24. doi: 10.1038/nn.2703.

[38] Schindelin J, Arganda-Carreras I, Frise E, Kaynig V, Longair M, Pietzsch T, Preibisch S, Rueden C, Saalfeld S, Schmid B, Tinevez JY, White DJ, Hartenstein V, Eliceiri K, Tomancak P, Cardona A. 2012. Fiji: an open-source platform for biological-image analysis. Nature Methods 9:676–682. doi:10.1038/nmeth.2019.

